# Enhanced specificity of *Bacillus* metataxonomics using a *tuf*-targeted amplicon sequencing approach

**DOI:** 10.1101/2023.05.28.542609

**Authors:** Xinming Xu, Lasse Johan Dyrbye Nielsen, Lijie Song, Gergely Maróti, Mikael Lenz Strube, Ákos T. Kovács

**Affiliations:** Bacterial Interactions and Evolution Group, DTU Bioengineering, Technical University of Denmark, 2800 Lyngby, Denmark; Institute of Biology Leiden, Leiden University, 2333 BE Leiden, The Netherlands; BGI-Tianjin, BGI-Shenzhen, 300308 Tianjin, China; Institute of Plant Biology, Biological Research Center, ELKH, 6726 Szeged, Hungary; Bacterial Ecophysiology and Biotechnology Group, DTU Bioengineering, Technical University of Denmark, 2800 Lyngby, Denmark

**Keywords:** Bacillus, amplicon sequencing, metataxonomics, tuf, species abundance

## Abstract

*Bacillus* species are ubiquitous in nature and have tremendous application potential in agriculture, medicine, and industry. However, the individual species of this genus vary widely in both ecological niches and functional phenotypes, which, hence, requires accurate classification of these bacteria when selecting them for specific purposes. Although analysis of the 16S gene has been widely used to disseminate the taxonomy of most bacterial species, this gene fails proper classification of *Bacillus* species. To circumvent this restriction, we designed novel primers and optimized them to allow exact species resolution of *Bacillus* species in both synthetic and natural communities using high-throughput amplicon sequencing. The primers designed for the *tuf* gene were not only specific for the *Bacillus* genus but also sufficiently discriminated species both *in silico* and *in vitro* in a mixture of 11 distinct *Bacillus* species. Investigating the primers using a natural soil sample, 13 dominant species were detected including *Bacillus badius*, *Bacillus velezensis*, and *Bacillus mycoides* as primary members, neither of which could be distinguished with 16S sequencing. In conclusion, a set of high-throughput primers were developed which allows unprecedented species-level identification of *Bacillus* species, including agriculturally important species.

## Introduction

The *Bacillus* genus is a prolific and diverse prokaryotic genus consisting of more than 100 species with validly published names, widely distributed in soil, sediment, air, marine environment, and even human systems [1, 2]. Members of the *Bacillus* genus comprise important species with economic, medical, and sustainability values as well as pathogenic strains. As an example, the *Bacillus cereus sensu lato (s.l.)* group includes the human pathogen *Bacillus anthracis*, the food poisoning agent *B. cereus*, and insect biopesticide *Bacillus thuringiensis* [3]. On the other hand, other members, such as *Bacillus subtilis*, *Bacillus amyloliquefaciens*, and *Bacillus velezensis* are widely used as biological control agents for both plants and animals due to traits such as spore-forming ability, high efficiency in plant root colonization, and abundant secondary metabolite production [4–7]. Additionally, *B. subtilis* has been a major cell and molecular biology model organism for decades [8]. This species has been extensively used for understanding bacteria biofilm formation, industrial production of enzymes and probiotics, and recently, as a proxy demonstrating phage-encoded biosynthetic gene clusters, and a non-photosynthetic bacteria that entrained circadian rhythm [9–11].

*Bacillus* as one of the most extensively studied plant growth promoting rhizobacteria (PGPR), it can competitively colonize plant roots and act as biofungicides, biofertilizers or biopesticides [12]. Despite the fact that members of *Bacillus* have been used as biological control agents for decades, classification and phylogenetic organization have been elusive for years [13, 14]. A current study demonstrated most *Bacillus spp.* registered as plant pathogen crop protection products have inconsistent species names in respect to current nomenclature [15]. One factor that caused the extensive polyphyly and misleading classification is the application of loose morphological criteria for assigning distinct species according to their cell shape and the ability to form spores [16]. Furthermore, as a conventional method to inspect the taxonomy of new isolates, members of the *Bacillus* genus commonly carry multiple copies of the 16S rRNA operon and these not only diverge widely within genomes, but also overlap extensively across different species, making it impossible to use 16S sequencing for species delineation [17]. Moreover, overall genetic differences are poorly correlated with phenotypic traits, and have even highlighted distinct phenotypes and genetic traits in strains with identical 16S alleles [18]. Given the wide variety of *Bacillus* species, specifically considering their role as both pathogens and biocontrol agents, it would appear urgent to develop novel approaches to alleviate the shortcomings of current methods.

To improve precise species-level identification of *Bacilli*, several alternative loci on the genome have been tested as phylogenetic discriminators of *Bacillus* species, including genes encoding gyrase subunit A (*gyrA*), the gyrase subunit B (*gyrB*), the RNA polymerase beta (*rpoB*), and elongation factor thermal unstable Tu (*tuf*) [19–28]. Caamaño-Antelo *et al.* isolated 20 foodborne *Bacillus* strains and analyzed the usefulness of three housekeeping genes, *tuf*, *gyrB*, and *rpoB* in terms of their discriminatory power. The *tuf* gene exhibited the highest interspecies similarities with sufficient conserved regions for primer matching across species whilst also containing enough variable regions for species differentiation [22]. Another study combined pulse field gel electrophoresis with *tuf* identification to successfully genotype *Bacillus* isolates from various environments [27]. Altogether, these studies suggested *tuf* as a potential phylogenetic marker among *Bacillus* species, and this gene has moreover been used for similar purposes in other genera [29].

In recent years, microbiologists have shifted their focus from single cultures to more complex microbial communities. This shift has come about partly as a reflection of the natural lifestyle of most bacteria, but more importantly, from observing widely different profiles of multi-cultured bacteria relative to their single-cultured counterparts. Although mostly single strain is included in biocontrol products for plants, natural environments have complex compositions containing diverse species of the *Bacillus* genus that may jointly function as plant growth promoters to alleviate abiotic stress or suppress plant pathogens [30–32]. In such a scenario, merely identifying individual taxa as *Bacillus* rather than individual species be insufficient for both basic science and potential microbiome engineering. Therefore, *Bacillus* community composition needs to be described in these complex settings. However, such analysis relies either on culture-dependent and laborious approaches or resource-extensive metagenomics which is unfeasible for high-throughput analysis. More commonly, amplicon sequencing is used to infer microbial community composition culture-independently and cost-efficiently [33]. Although amplicon analysis of the 16S rRNA gene has been remarkably successful owing to universality in bacteria, this method does not allow confident species identification not only in *Bacillus* genus but has low resolvability in other medically important genera, e.g. *Staphylococcus* and *Pseudomonas*. Thus, alternative molecular markers including *cpn60*, *rpoB*, and *gyrB* have been developed for universal bacterial genotypic identification, and whilst these genes have high discriminatory power, they lack universally conserved regions for primers and correspondingly for amplicon analysis [34–36]. While these approaches offer greater resolution during bacterial community profiling, they remain taxa specific. For instance, *gyrB* reveals highly similar bacterial community structure within Proteobacteria and Actinobacteria when compared with 16S, but it has merely 21.5% consistency in Firmicutes [34]. Therefore, amplicon sequence tools have been already developed for few genera that enable species-level resolution, such as *rpoD* amplicon methods for *Pseudomonas* and *tuf*-based methodology for *Staphylococcus* genus [29, 37].

Since differentiation of *Bacillus* in mixed communities is highly relevant, but impossible with standard 16S sequencing, we developed primers for species level differentiation. Specifically, we investigate conserved genes and their corresponding primers for *Bacillus* species identification using an *in-silico* amplification approach using *Bacillus* versus non-*Bacillus* genomes. Subsequently, *tuf* gene specific primer pairs were designed and inspected leading to a primer pair with high accuracy and specificity on *Bacillus* species. Finally, an amplicon sequencing method was tested on an Illumina MiSeq PE300 platform based on the selected primers, along with a customized database for *Bacillus* taxonomic assignment. The identified *tuf2* primers demonstrated almost full coverage of *Bacillus* species along with discriminatory power approaching whole genome phylogeny. Moreover, the *tuf2*-based amplicon approach allowed *Bacillus* profiling in natural communities, which we believe will facilitate large-scale exploration of bioactive *Bacillus* species or keep track of activeness of bio-inoculants.

### Material and methods

### Primer design

In response to the insufficient availability of differentiating primer pairs for *Bacillus* genus, an exhaustive search for *Bacillus* conserved genes was conducted to uncover genes with high phylogenetic discrimination power as candidate targets for primer design. We chose housekeeping genes that were frequently employed in literature and prioritized *rpoB*, *gyrA*, and *tuf* as our candidate genes to design primers. To evaluate the breadth of coverage for these primers, 1149 complete genome sequences of *Bacillus* were downloaded using *ncbi-genome-download* with RibDif and added to our *Bacillus* genome collection (listed in Dataset S1). A few *Bacillus* genomes were re-identified with TYGS due to the poor annotations of uploaded genomes to improve the phylogenetic inference accuracy. Pan-genome analyses were carried out using roary, showing these three genes (*rpoB*, *gyrA*, and *tuf)* are indeed core genes (presented in >99% of the strains in our *Bacillus* genome collection) [38]. Meanwhile, previously documented primers targeted on these genes have high amplification rate against the *Bacillus* genome.

Next, sequences of candidate genes were then dereplicated with vsearch followed by sequence alignments using MUSCLE v5 [39, 40]. Analysis of multiple sequence alignments (MSAs) was conducted to target conserved regions flanking highly variable regions of 300-600bp (https://github.com/mikaells/MSA-primers). Potential primers were suggested from these conserved regions. Out of several preliminary primer designed, the primer pair *tuf1-F* (5’-CACGTTGACCAYGGTAAAACH-3’), *tuf1-R* (5’-DGCTTTHARDGCAGADCCBTT-3’) and *tuf2-F* (5’-AVGGHTCTGCHYTDAAAGC-3’), *tuf2-R* (5’-GTDAYRTCHGWWGTACGGA-3’) targeting a 500 bp sequence of *tuf* gene showed the top performance characteristic in initial examination and was selected for further evaluation.

### *In silico* evaluation of primers

Initially, RibDif was used to evaluate the usefulness of standard primers for taxonomy, which specifically means primers targeting the V3V4 and V1V9 region of the 16S gene in bacteria [17]. As this analysis clearly showed the inability of these standard primers in terms of resolving the individual species of *Bacillus*, we investigated 11 primer sets targeting the genes of 16S rRNA, *gyrB*, *gyrA*, and *rpoB* derived from previous studies along with our two newly designed sets of *tuf* primers. We used RibDif to download all completed genomes of the *Bacillus* genus and then evaluated the performance of these primers on this collection through *in silico PCR* (https://github.com/egonozer/in_silico_pcr).

We followed the standards as described by *Lauritsen et al.* to benchmark the performance of primers according to two metrics, both of which should encapsulate the functional performance in mixed communities [37]. Specifically, 1) what is the proportion of *Bacillus* genome amplified and 2) what is the proportion of non-*Bacillus* genome amplified. Furthermore, the top candidate primers were further evaluated by building phylogenetic trees from the alignments of their resulting amplicons. These phylogenetic trees were inferred using neighbor-joining (NJ) method with the Maximum Composite Likelihood model, and 1000 bootstraps where used to test the strengths of the internal branches of the trees. Trees were visualized in iTol (https://itol.embl.de/). TreeCluster were used to define clusters within the trees and these clusters where then compared to the known taxonomy of the amplicons [41]. Cohens Kappa was calculated with the R package “irr” to infer the degree of agreement between TreeCluster and the known species names [42, 43].

### Whole genome sequencing

A short and long read hybrid approach was used to sequence new *Bacillus* isolates obtained from an ongoing project in our laboratory. Bacterial genomic DNA was extracted using E.Z.N.A.® Bacterial DNA Kit (Omega, Bio-tek, USA, Georgia). The qualities and quantities were evaluated by NanoDrop DS-11+ Spectrophotometer (Saveen Werner, Sweden, Limhamn) and Qubit 2.0 Fluorometer (Thermo Fisher Scientific). The libraries for short-reads sequencing were constructed using the MGI paired-end protocol [44]. Briefly, 300ng DNA was fragmented to 200-300 bp using segmentase enzyme followed by fragment selection with VAHTS™ DNA Clean Beads (Vazyme; China, Nanjing). Subsequently, end repair, A-tailing reactions and adapter ligation were implemented. After PCR and purification, the libraries were sequenced on the MGISEQ-2000 (MGI Tech Co., Ltd.) platform according to the manufacturer’s instructions to generate 2× 150 bps paired end reads. For Nanopore sequencing, the rapid barcoding kit (SQK-RBK110.96) was used and these libraries were sequenced with an R9.4.1 flow cell on a MinION device running a 48-h sequencing cycle. The resulting reads were base called and demultiplexed with MinKNOW UI v.4.1.22. For *de novo* assembly, the NGS short reads were adapter and quality trimmed using fastp v.0.22.0 and the Nanopore reads were adapter trimmed using porechop v.0.2.1 [45, 46]. The trimmed reads from Nanopore were assembled using flye v.2.9.1-b1780, and subsequently the trimmed reads from both platforms and the long read assembly were hybrid assembled with Unicycler v.0.5.0 using the *–existing_long_read_assembly* option [47, 48]. The completeness and contamination levels of each strain was checked using CheckM v.1.2.2 [49]. The assemblies were then taxonomically assigned and placed in the full-genome, multi locus GTDB-Tk reference tree, using the Classify Workflow of GTDB-Tk v2.1.1 [50]. The tree was subsequently pruned to create a full-genome multi locus tree of the query strains. The chromosomes were annotated using the NCBI Prokaryotic Genome Annotation Pipeline [51].

### Comparative identification of *Bacillus* isolates

Since the primers targeting the *tuf* gene appeared superior for *Bacillus* differentiation, we investigated these in detail. Initially, we assessed the accuracy of the *tuf* primers by comparing the resolving power with other means of *Bacillus* identification methods, such as 16S rRNA PCR and whole genome sequencing. Complete 16S rRNA genes were PCR-amplified with primer pair 27F (5’-AGAGTTTGATCCTGGCTCAG-3’) and 1492R (5’-GGTTACCTTGTTACGACTT-3’) [52]. A 25 µl PCR mixture contains: 2.5 µl 2 mM dNTP, 2.5 µl 10× DreamTaq Buffer, 0.5 µl of each primer (10 µmol/l), 0.25 µl DreamTaq DNA Polymerase (5 U/µl), 17.5 µl nuclease-free water, and 1.25 µl DNA template. Thermal cycling conditions were 95 °C for 3 min; 30 cycles of 30 s denaturation at 95 °C, 30 s annealing at 55 °C, and 1 min extension at 72 °C; final extension at 72 °C for 10 min. PCR product were purified using NucleoSpin gel and PCR cleanup kit (Macherey-Nagel; Germany, Düren) and sent for Sanger sequencing Eurofins Genomics.

### Evaluation of primer performance on amplicon sequencing

We chose 11 distinct *Bacillus* strains to create a synthetic community to evaluate the *in vitro* performance of the *tuf* primers on species resolution through amplicon sequencing. *Bacillus thuringiensis* 407 cry^-^ (NCBI accession number GCF_000306745.1), *Bacillus velezensis* SQR9 (CGMCC accession number 5808), *Bacillus cereus* ATCC 14579 (NCBI accession number GCF_000007825.1), and *Bacillus subtilis* PS216 (NCBI accession number GCF_000385985.1) were type culture collection strains [53–56]. The rest seven isolates were obtained from ongoing projects in our laboratory and identified by whole genome sequencing. All bacteria were grown in lysogeny broth (LB; Lennox, Carl Roth, Germany, Limhamn) overnight, and supplemented with 28% glycerol before storing them at −80 °C. DNA extractions of each strain were pooled in equimolar ratio to create a positive control mixture (Bac-DNAmix). To benchmark the performance of *tuf2* on amplicon sequencing, primer pairs *gyrA3* that were previously applied on *Bacillus* mock community (*gyrA3-F*: 5′-GCDGCHGCNATGCGTTAYAC-3′ and *gyrA3-R*: 5′-ACAAGMTCWGCKATTTTTTC-3′) and universal primers targeting the V3-V4 hypervariable region of the 16S rRNA gene (341F: 5’-CCTACGGGNGGCWGCAG-3’ and 805R: 5’-GACTACHVGGGTATCTAATCC-3’) were selected [20, 52]. Short barcodes were attached on all primers for downstream sequence demultiplication as is listed in Table S1. For *tuf2* amplification, a 25 µl PCR mixture contains: 12.5 µl TEMPase Hot Start 2× Master Mix Blue, 0.8 µl of each primer (10 µmol L^-1^), 10.6 µl nuclease-free water and 0.3 µl DNA template. The PCR program included initial denaturation for 15 min at 95 °C; 30 cycles of 30 s at 95 °C, 30 s at 47 °C and 1 min at 72 °C; and a final extension for 5 min at 72 °C. For the amplification of 16S rRNA and *gyrA*, the annealing temperatures were 62 °C and 50 °C, respectively. All PCR products were purified and pooled into equimolar ratios and sequenced on a MiSeq platform using the MiSeq Reagent Kit v3(600-cycle).

### Amplicon data analysis

Raw sequence data was processed with the QIIME2 pipeline for all primer sets [57]. Primers and barcodes were removed with cutadapt, and after demultiplexing, amplicons were denoised, merged, and chimera-checked using DADA2 [57, 58]. In *Bacillus* DNA mixture, all ASVs were assigned to NCBI database for parallel comparison to avoid bias. For natural soil sample, 16S data were analyzed using standard workflow SILVA database and Naive Bayers classifier [59]. A *tuf*-specific database was built from all *tuf* genes of our *Bacillus* genome collection (available at https://github.com/Xinming9606/BAST). The *tuf* amplicons of each sample where then taxonomically assigned using BLASTN against the *tuf-*database, using *max_target_seq: 1*. was used to assign taxonomy to the representative sequences of amplicon data with the best hits selected as taxonomy names.

### Data availability

The raw sequencing data has been deposited to NCBI Sequence Read Archive (SRA) database under BioProject accession number PRJNA960711 and PRJNA976106. All code is available at https://github.com/Xinming9606/BAST.

## Results

### Comparative analysis of primer pairs for *Bacillus* identification

A battery of six genes and primers widely used for *Bacillus* identification were compared using *in silico PCR*. Primers were tested against a *Bacillus* genome collection including 1149 complete genomes downloaded from NCBI in April 2023, along with 41 non-*Bacillus* genomes corresponding to other well-studied microbes often found in soil (listed in Dataset S1 and Table S2).

Initially, the performance of the normally used primers targeting the 16S gene was examined to provide a baseline and motivation for our investigation. As they are designed for, the universal primer sets targeting the full (V1V9) and partial (V3V4) parts of the 16S rRNA gene successfully amplified both *Bacillus* and non-*Bacillus* strains (Table 1). Using the RibDif tool for a more detailed analysis, however, revealed that *Bacillus* has an exceptionally high allele multiplicity and extensive species overlap, which means that amplicons derived from 16S rRNA gene will rarely be unique for individual species. For example, full-length V1V9 amplicons derived from *B. subtilis* are indistinguishable from amplicons derived from *B. velezensis*, *B. siamensis* and *B. amyloliquefaciens*. Thus, 16S gene it is not an ideal molecular marker for the *Bacillus* genus [18]. The *B. subtilis* group specific primers Bsub5F and Bsub3R reportedly can identify *B. subtilis* group exclusively, including species *B. subtilis*, *Bacillus pumilus*, *Bacillus atrophaeu*s, *Bacillus licheniformis* and *B. amyloliquefaciens.* This primer set had hits on 797 *Bacillus* genomes out of the 1149 (69.36%), and did not amplify non-*Bacillus* genomes. Despite a high level of specificity, the lack of broad coverage in these amplified sequences will not provide species-level identification in diverse communities.

**Table 1.**
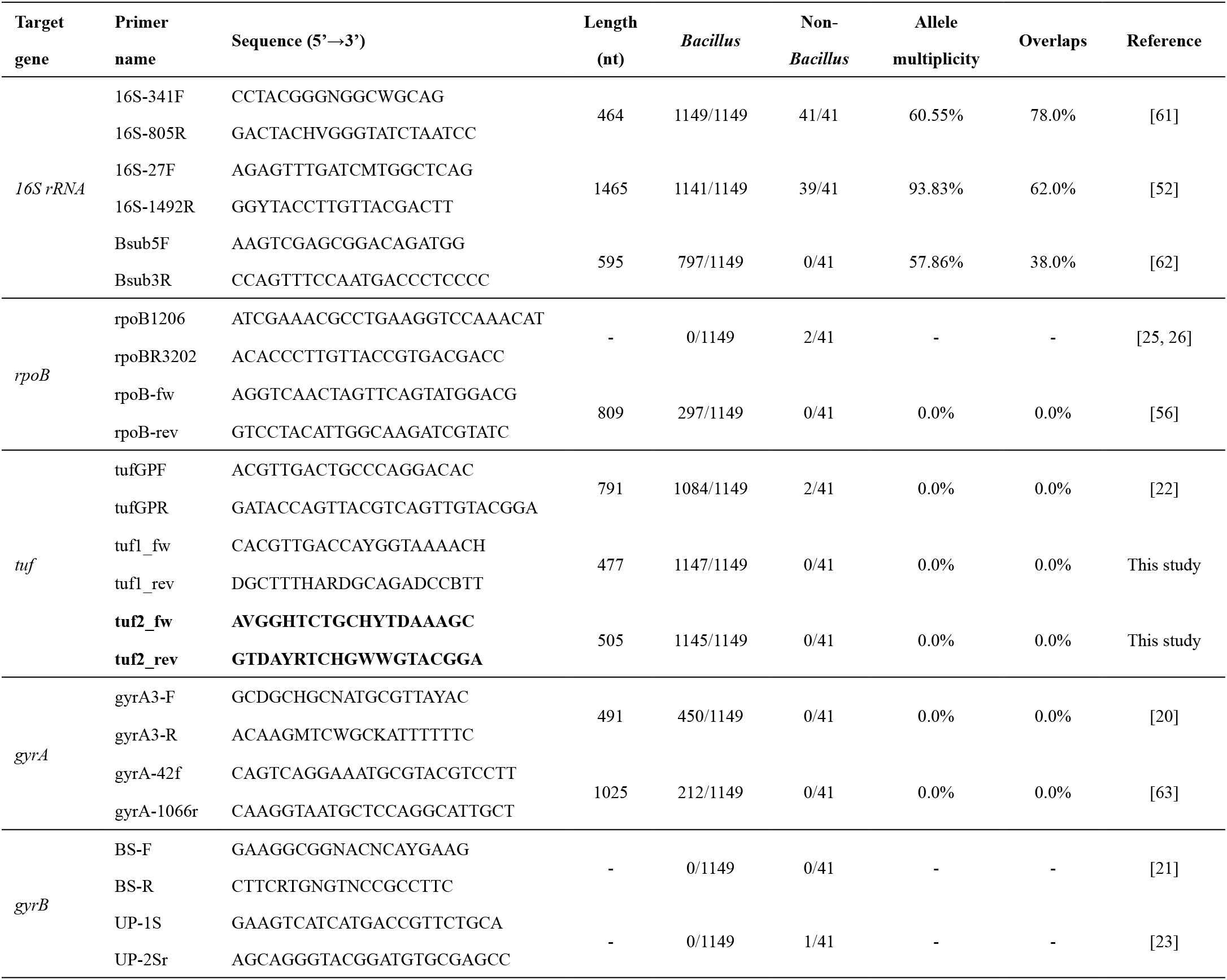
Summary of published and newly developed *Bacillus* identification marker genes and corresponding primer pairs. The amplification rates of *Bacillus* genomes represent the universality of each primer on various species of *Bacillus*, and the amplification rates on non-*Bacillus* genomes show the specificity of primer pairs. Allele multiplicity is the percentage of genomes having more than one unique allele and overlap is the percentage of species having at least one allele overlapping another species.

Next, we investigate primer pairs targeting housekeeping genes. Previous studies reported that the partial sequence of gyrase subunit A sharply separated twelve strains belonging to *B. amyloliquefaciens*, *B. atrophaeus*, *B. licheniformis*, *B. mojavensis*, *B. subtilis* and *B. vallismortis* [60]. Liu and colleagues have also reported their primer pair *gyrA3* distinguished 6 species in the mock community they constructed. During our examination, the *gyrA*-based primer pairs displayed high specificity for the *Bacillus* genus with no amplification of species from the other genera. However, the primers targeting the *gyrA* gene amplified only 39% (450/1149) and 18% (212/1149) of the genomes, using *gyrA3-F - gyrA3-R* and the *gyrA*-42f - *gyrA*-1066r primer combinations, respectively, suggesting lack of discriminatory potential among certain clades within the *Bacillus* genus.

Similarly, the primer pair targeting a 809-nucleotide region of the *rpoB* gene displayed limited detection with 26% (297/1149) amplification rate by primers *rpoB-fw* and *rpoB-rev*. Surprisingly, some of the published primer sets which claimed to drastically increase resolution and/or discrimination displayed no amplification of *Bacillus* genomes at all, which was especially evident for *gyrB*-based primers, such as the *BS-F - BS-R* and the *UP-1S - UP-2Sr* sets that were designed for *B. cereus* group identification. In contrast, the primer pair tufGPF and tufGPR, targeting the *tuf* gene displayed high amplification rate, although the amplicon generated (791 bp) are longer than the Illumina technology currently supports for overlap, and the resulting sequencing gap complicates analysis and may decrease the resolution of species identification.

Motivated by the lack of generally applicable and sufficiently differentiating primer pairs for the *Bacillus* genus, we designed primer pairs targeted on *tuf* gene. Two sets were investigated, since 4 conserved regions within the *tuf* gene alignment were available (position 58, 517 to 518, and 958 to 1004. Thus, primer pairs where likely to be at 58 to 517 and position 517 to 1004 (Fig. S1B). Profiling the *tuf* gene for nucleotide diversity showed that these conserved regions flanked variable regions of high nucleotide diversity which may potentially allow species identification (Fig. S1A). In total, 18 *tuf*-based primer pairs were suggested (Dataset S2). According to the lowest number of degenerate sites and substitutions within the primer sequences, two sets of primers were selected for further evaluation. Both of these primers, now referred to as *tuf1* and *tuf2*, had substantially better performances than the ones hitherto tested: it had the highest coverage at close to 100% amplification rate, along with a notable 0% rate of non-*Bacillus* amplification from the genomes of the negative controls. From these results, both primer sets targeting the *tuf* gene were selected for further analysis.

Of note, the predicted primer pairs for *rpoB* and *gyrA* loci had high number of degenerate sites and the most conserved fragments in the alignments remained highly diverse. For example, only two primer pairs between position 1549 and position 1927 were suggested for *rpoB* alignment, with more than 10 sequences incorporating three degenerate nucleotides (Fig. S2). Thus, primer design is challenging on such non-conserved fragments.

### Phylogenetic analysis of *Bacillus* species based on different sequence approaches

To evaluate the discrimination resolution of the selected *tuf* primers, phylogenetic trees of their *in silico* PCR amplicons alignments were created for both along with the amplicons generated with 16S V3-V4 primers as comparison (Fig. 1). Two approaches were applied to analyze the phylotaxonomic distribution obtained using the *tuf1, tuf2* and 16S gene amplicons: neighbor-joining trees were used to show the phylogenetic distribution of the amplicons and the TreeCluster program was then used to cluster the amplicons on the basis of their position in these trees. The reasoning for this was to compare the phylogenetic positioning with known taxonomy, which consequently reveals how well these amplicons can group their parent genome correctly. The phylogenetic tree of *tuf2* amplicons grouped the amplicons in correspondence with the published species names of their parent genomes. Divergent clades were interspersed by different species with the main division of subtilis clade and cereus clade. Few branches displayed inadequate separation of nodes, although this was mainly due to poor annotation or species misnaming, e.g, certain *B. velezensis* isolates were originally proposed as *B. amyloliquefaciens*. The inter-rater reliability analysis using Cohen’s kappa showed a substantial agreement (kappa = 0.721, z = 35.6, p < 0.001) between known species names and *tuf2* amplicon clusters. The tree of the *tuf1* amplicons performed substantially worse in this regard, having much less systematic clustering of each taxa (kappa = 0.326, z = 21.9, p < 0.001). Thus, we do not recommend *tuf1* primers for *Bacillus* identification. As expected, the tree based on 16S rRNA genes exhibited worse intraspecific phylogenetic resolution than *tuf2*, such as failing to delineate the distinct groups within *B. cereus*, *B. anthracis*, and *B. thuringiensis*. Moreover, TreeCluster annotation revealed distinct species having identical 16S V3-V4 sequences as well as multiple instances of one genome having 16S alleles in several clades of the tree (kappa = 0.63, z = 26.9, p < 0.001). Noteworthy, 60% of *Bacillus* genomes exhibit multiple alleles and V3V4 sequences were extensively dereplicated for tree construction.

**FIG. 1.**
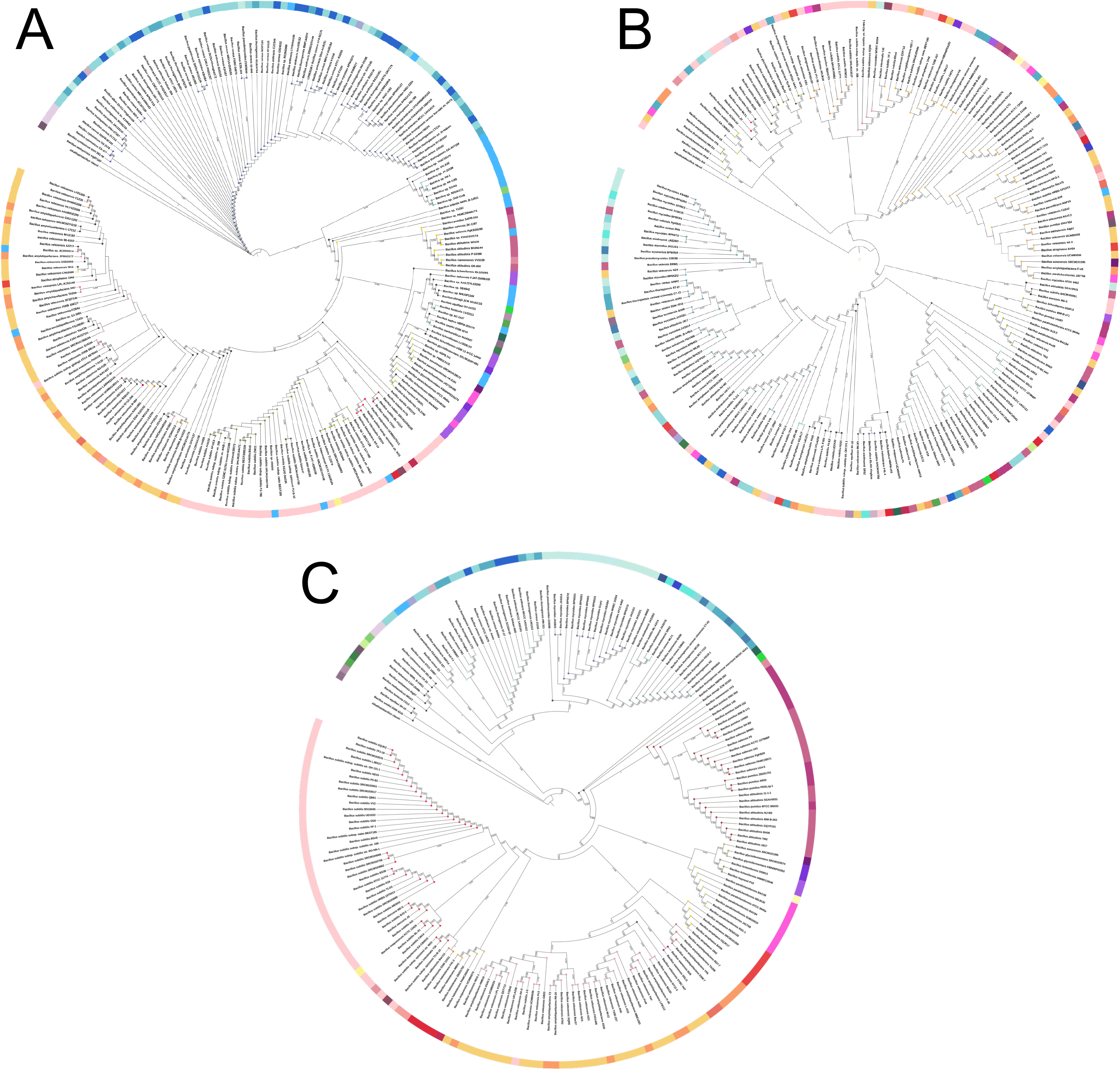
Phylogenetic tree of the *in-silico* PCR product using **A** 16S rRNA, **B** *tuf1*, **C** *tuf2* primer pairs. Species names were annotated by whole genome sequencing. The outer ring was colored based on published species names and circles on the tips denoted these species have identical amplicons. *Alkalihalobacillus clausii* was used as the outgroup to root the tree.

To validate the accuracy of the *tuf2* primer pairs *in vitro*, we compared the phylogenetic affiliation between amplicons of 16S rRNA, *tuf2*, and complete genomes of twenty soil-derived isolates (Fig. 2). The same genomic DNAs were used for Sanger sequencing and whole genome sequencing. The *tuf2*-based tree clearly delineated seven distinct clusters with high bootstrap values and in good agreement with the tree structure depicted based on the whole genomes. For instance, *tuf2*-tree grouped environmental strains, D8_B_37, G1S1 etc. in the *Peribacillus* cluster as expected since the sequence identity with *Peribacillus simplex* was >98% shown on NCBI-blast (Table S3). Similarly, isolates were accurately grouped that belong to *B. subtilis*, *B. licheniformis*, *B. velezensis*, although it demonstrated lower resolution on the identification of *Bacillus altitudinis*, *B. pumilus* and *B. safensis*. The 16S rRNA gene-based tree lacked concordance with full genome-based phylotaxonomics and led to an unreliable phylogenetic signal due to the prevalence of multiple copies of 16S in *Bacilli* in addition to the high genetic similarity of 16S rRNA genes between these species. It is worth mentioning that *in vitro* assay *tuf2* primers did not exhibit any unspecific amplification of negative control *Pseudomonas aeruginosa, Streptomyces iranensis, and Clavibacter michiganensis* in line with expectations from *in silico* tests (Fig. S3).

**FIG. 2.**
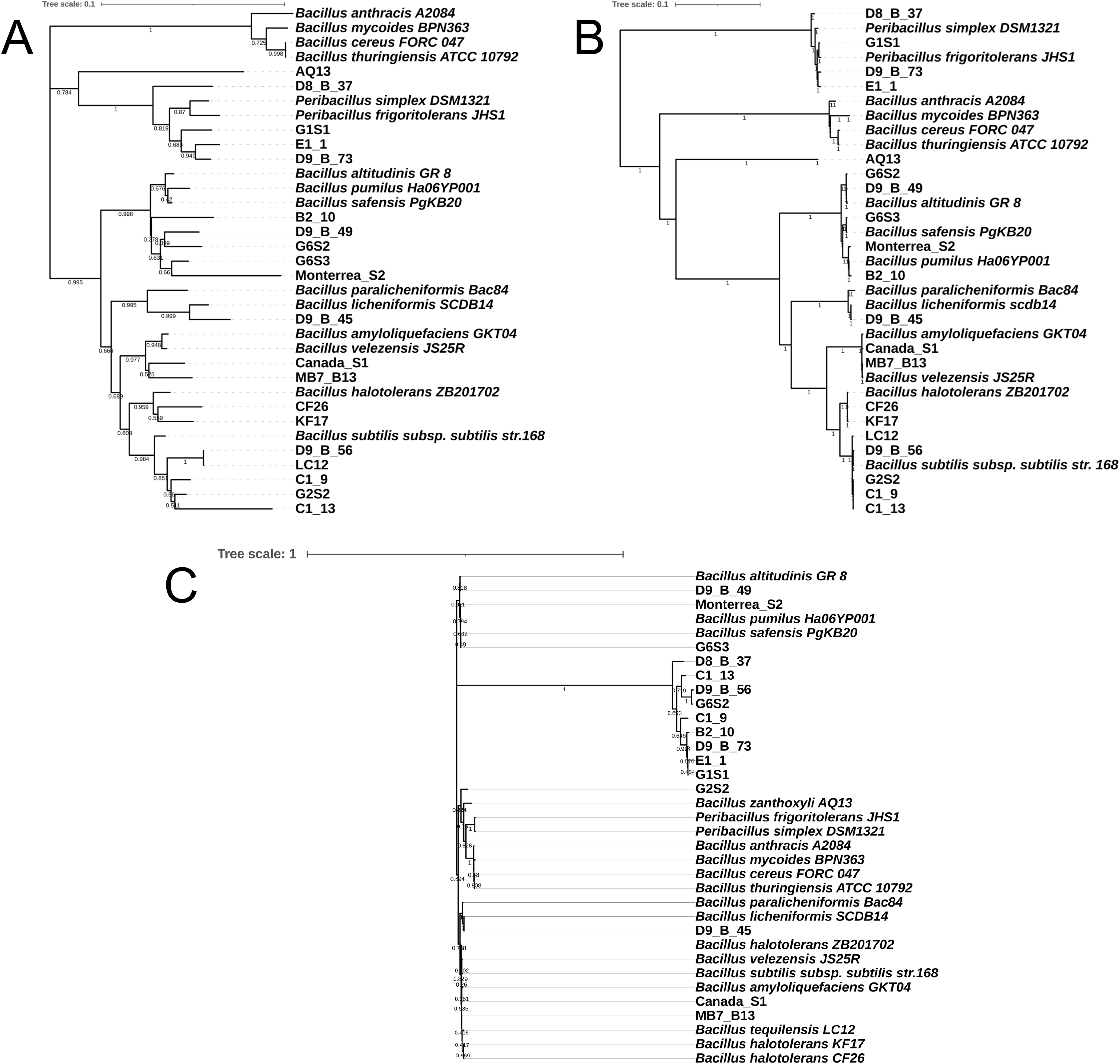
Phylogenetic analysis of (A) amplified *tuf* gene, (B) complete genome, and (C) 16S rRNA gene of *Bacillus* spp. that were recently isolated or strains with publicly available genomes in NCBI GenBank was obtained using the Neighbour-Joining method. Numbers depicted on the branches indicate bootstrap values.

In summary, the *tuf2* primers developed in this study is an effective tool to identify the species-level taxonomy within the *Bacillus* genus with high phylogenetic discrimination power comparable to the methods based on complete genome.

### Amplicon Sequencing of *Bacillus* Synthetic Community

Next, we investigated whether the *tuf2* primer pairs could be applied for high-throughput amplicon sequencing. To evaluate the specificity and efficiency of *tuf2*, we compared these with the frequently employed 16S rRNA primers (V3-V4) and a newly published primer set (*gyrA3*) that was suggested to have the potential for Illumina sequencing of complex *Bacillus* community [20, 52]. A defined DNA mixture containing 11 *Bacillus* species was assembled and sequenced by the three sets of primers.

As expected, 16S V3V4 amplicon sequencing performed poorly, only identifying 4 out of 11 species and instead overestimating species in the *B. subtilis* group or the *B. cereus* group, resulting in *B. cereus* abundance being highly overinflated. Within the subtilis clade, *B. velezensis* was three-fold larger than expected. Apart from the 16S gene-based identification only provided correct detection of 4 species, this method instead inaccurately reported *Priestia aryabhattai* within the sample composition which was not added to the DNA mixture (Fig. 3). The approach with the primer pairs of *gyrA3* locus resolved 8 species, including *B. altitudinis*, *B. licheniformis*, *B. velezensis* that were previously validated during development of these primer pair, but largely overestimated the proportion of *B. pumilus* and *B. safensis* (Table 2). In comparison, the *tuf2* amplicon method was able to identify 9 strains but missed *B. amyloliquefaciens* and *B. cereus*, other species have closer correspondence to expected abundance and lower deviation. These data suggest not only that *tuf* specific primers can reveal molecular variation at species level, but also complements and potentially outperforms 16S rRNA amplicon sequencing in complex *Bacillus* community studies. Noteworthy, all amplicons were assigned to NCBI database for parallel comparison which includes partial sequences and incorrect information, thus, mapping to our customized *tuf* database would highly improve the accurateness of results.

**FIG. 3.**
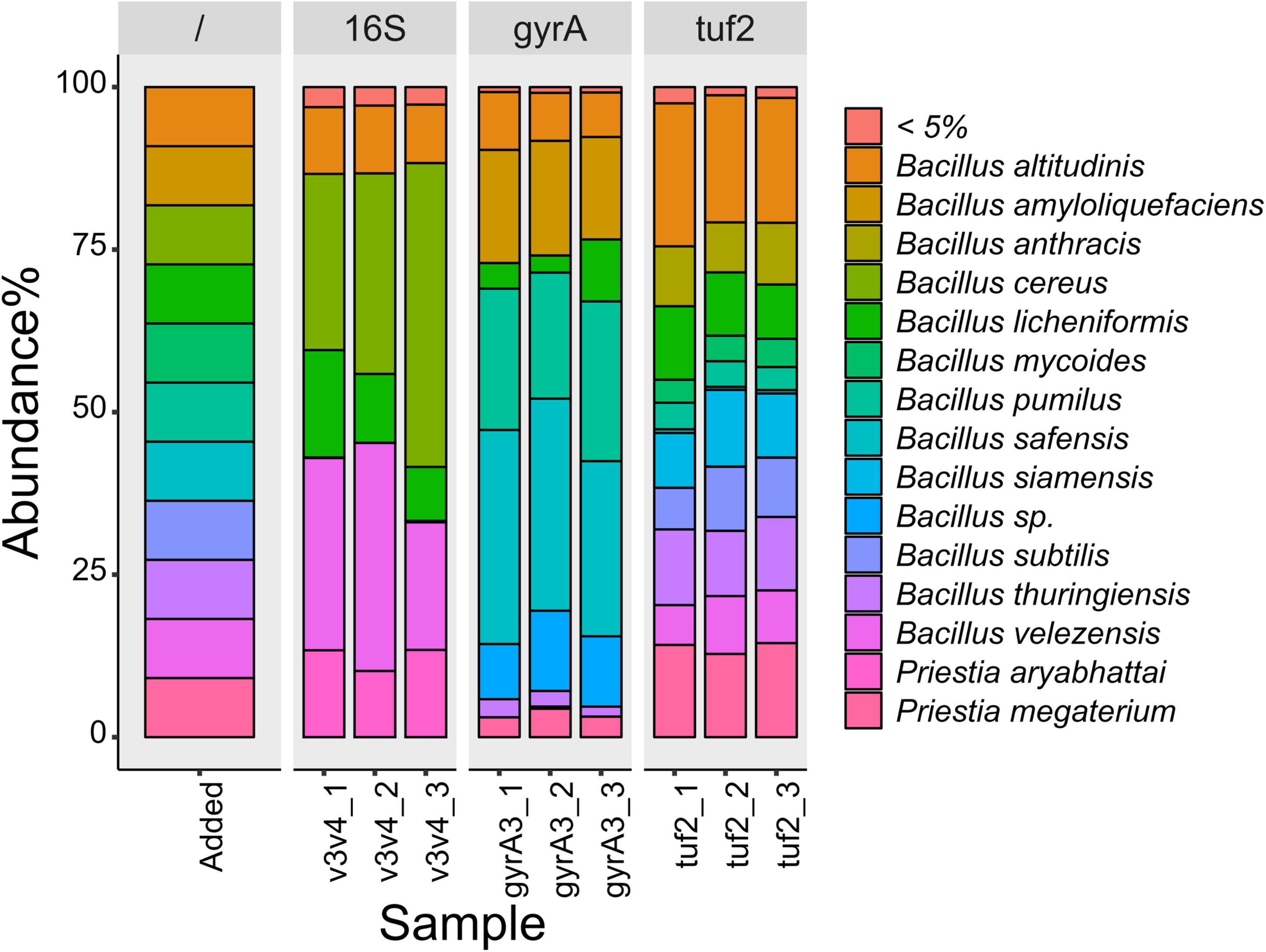
Relative abundance of each species in the Bac-DNAmix containing the mixture of 11 *Bacillus* species. The first bar represents the theoretical abundances in the *Bacillus* DNA mixture followed by abundances detected using the V3-V4 *16S rRNA*, *gyrA*, and *tuf* 2 sequencing approaches, respectively.

**Table 2.** Composition of Bac-DNAmix revealed by four primer sets.

| Species name | Strain | 16S relative abundance<br>(%)* | *gyrA3* relative<br>abundance (%) | *tuf2* relative abundance<br>(%) |
| --- | --- | --- | --- | --- |
| *B. altitudinis* | G6S2 | 108.89 ± 7.01 | 84.75 ± 9.60 | 222.77 ± 13.63 |
| *B. amyloliquefaciens* | B12 | 0 | 186.00 ± 9.16 | 0 |
| *B. cereus* | ATCC14579 | 383.84 ± 93.61 | 0 | 0 |
| *B. licheniformis* | D9_B_45 | 129.81 ± 38.08 | 58.93 ± 32.96 | 107.72 ± 13.26 |
| *B. mycoides* | SIN1.1 | 0 | 0 | 43.33 ± 3.68 |
| *B. pumilus* | Monterrea_S2 | 0 | 240.93 ± 23.27 | 42.38 ± 2.54 |
| *B. safensis* | G6S3 | 0.76 ± 0.43 | 339.09 ± 30.29 | 5.70 ± 0.47 |
| *B. subtilis* | PS216 | 0 | 0 | 93.03 ± 16.36 |
| *B. thuringiensis* | 407 | 0 | 24.73 ± 5.74 | 120.93 ± 7.61 |
| *B. velezensis* | SQR9 | 309.08 ± 70.36 | 1.19 ± 1.69 | 84.77 ± 12.88 |
| *Priestia megaterium* | B10 | 0 | 38.82 ± 6.62 | 152.06 ± 8.00 |

\*Estimation values for *tuf* relative abundance versus theoretical abundance are given as mean ± standard deviation, where 100% is the expected value.

### Profiling *Bacillus* species in natural soil sample

To elucidate the feasibility of the newly developed primer pairs to dissect the composition of the *Bacillus* genus in natural soil samples, we used the *tuf2* primers, since they performed best in our analysis, and compared it with V3V4 sequencing. Using V3-V4 16S rRNA gene-based amplification, 53,517 reads were mapped for classification per sample on average, from which 99% of the reads were unidentifiable on the species level. A total of 741 families were found, with the dominant families belonging to *Xanthobacteraceae* (9%), *Chthoniobacteraceae* (3.2%), *Isosphaeraceae* (3.8%), *Bacillaceae* (6.1%). Out of 70 *Bacillus* ASVs, only 2 were annotated as *Bacillus sp.* and *B. simplex* in the environmental samples. It is conceivable to improve the species designation by assigning the sequence reads a defined *Bacillus* database, however the issue of heterogeneity and multiplicity of the 16S rRNA gene still remain. For *tuf2* sequencing, after filtering, denoising and chimera removing, an average of 28,231 reads per sample were available. Rarefaction curve showed saturated sequencing depth for all samples with 601 ASVs assigned (Fig. S4). With these primers, species level composition could be achieved, showing the predominant species in this soil sample to be *B. badius*, *Bacillus dafuensis*, *Bacillus infantis*, and *Bacillus weihaiensis*, which presumably can be considered the correct composition of the fraction classified as *Bacillus* in the 16S analysis (Fig 4). *B. velezensis*, *B. mycoides*, and *B. amyloliquefaciens* were also detected in natural soil with lower abundance. Our results demonstrated the ability of *tuf2* as a complementary analysis when employing 16S analysis for specifically profiling *Bacillus* species in natural soil. As an example, one would infer that since ∼10% of *Bacillus* is *B. velezensis* and ∼5% of overall bacteria is *Bacillus*, the total abundance of *B. velezensis* is ∼ 0.5%.

**FIG. 4.**
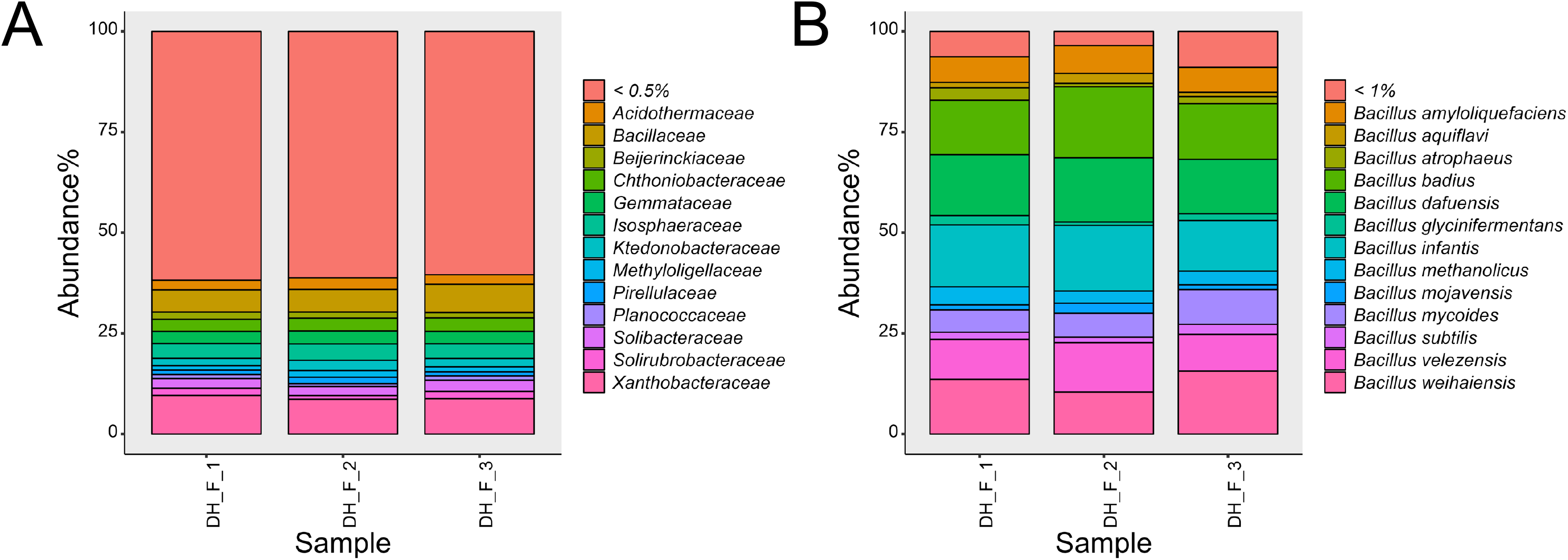
**A** Relative abundance of the most abundant bacterial families, and **B** Relative abundance of the most abundant *Bacillus* species in natural soil sample collected from Dyrehaven (55.791000, 12.566300).

## Discussion

The *Bacillus* genus is one of the most predominant bacterial genera in soil and exerts multiple direct and indirect effects beneficial for plants, including nitrogen fixation, nutrients acquisition, production of phytohormones, and anti-pathogenic effects [64, 65]. Nevertheless, in nature, rather than existing as a solitary microbiome comprising a single species, *Bacillus* spp. exist as a part of complex microbial community and therefore may serve as PGPRs by diverse species. The urgent need to switch to sustainable agriculture has boosted the demand for employing biological agents rather than chemical ones, and members of the *Bacillus* genus has come into focus as a joint PGPR group. Not all *Bacillus* are equally useful as PGPR agents, however, which means that methods that can accurately identify *Bacillus* spp. and systematically profile *Bacillus* communities in a natural soil has now become of vital importance. Here, we designed and scrutinized primer pairs that can dissect the *Bacillus* genus on species level. Moreover, a corresponding bioinformatic pipeline has been employed that allows simple analysis of Illumina Miseq300 platform-based data on QIIME2 enabling rapid identification and selection of soils with specific *Bacillus* communities for further analysis and culturing.

Previous studies have designed and used *Bacillus*-specific primer sets targeting non-universal regions of the 16S rRNA gene for rapid taxonomic identification, and alternative biomarkers *rpoB*, *gyrB*, and *gyrA* have been proposed to resolve the limited intra-specificity as well. For genotyping of single strains, collections of housekeeping genes, such as *gyrB* and *rpoB* are often used, e.g. in multilocus sequence analysis (MLSA) with different schemes targeting on different *Bacillus* groups. For instance, studies constructed phylogeny based on concatenated housekeeping genes *gyrB*-*rpoB*-*pycA*-*pyrE*-*mutL*-*aroE*-*trpB* to distinguish the *B. pumilus* group [66, 67]. However, use of these genes as universal markers in *Bacillus* performed poorly, as evident by *gyrB* and *rpoB* primer sets tested in our study had no amplification against the *Bacillus* genome collection [21, 23, 24, 68]. The lack of *in silico* amplification might potentially be caused by the lack of adjustment in annealing temperature and internal walking primers were not introduced in our test. Primer sets *gyrA-42f* and *gyrA-1066r* limited within *B. subtilis* group amplification but would still provide accurate classification and works for single isolate identification.

Short *tuf* gene sequences have been reported as a reliable molecular marker for investigating the evolutionary distances between *Lactobacillus* and *Bifidobacterium* [69]. Additionally, *tuf* gene sequencing has previously been employed for identification of *Staphylococcus* species in both single isolate and in a high-throughput context [29, 70]. Caamaño-Antelo compared interspecies sequence similarities among 20 *Bacillus* isolates using 3 loci and demonstrated 57,80%, 67,23%, and 77,66% similarities for *gyrB*, *rpoB*, and *tuf* gene alignments, respectively. Our results demonstrated that *tuf*-based primers had superior performance on the specificity and range, evident by the 100% amplification rate of 1149 *Bacillus* genomes and no unspecific amplification of negative controls. When using our *tuf* primers for the identification of soil isolates, Sanger sequencing aligned exactly with the results we obtained from the complete genome demonstrating the versatile use of *tuf* primers.

An important reason that 16S rRNA gene gained widespread use is the universality in bacteria combined with highly conserved regions that facilitate universal primer targets flanked by variable regions that are suited for metataxonomics on next-generation sequencing platforms. However, based on previous analysis conducted by RibDif, it has been found out that most genomes of *Bacillus* have multiple alleles of V3V4 region, and 39 out of 50 species have V3V4 alleles that are not unique to particular species [17]. As per RibDif analysis, a community containing *B. subtilis* analyzed with V3V4 metaxonomics will also incorrectly suggest several unique ASVs due to the multiple alleles of *B. subtilis* and hence overestimate the richness the sample. In a sample containing *B. thurigiensis*, one may even incorrectly infer the presence of no less than 14 other species, as all these have V3V4 alleles shared between one another. As a result, amplicons of the V3V4 region of the 16S rRNA gene cannot be used to differentiate species of *Bacillus.* Therefore, we aimed to generate primers which can separate species of *Bacillus* phylogenetically. When designing primers that targeted the *tuf* gene, consensus sequences were identified at the beginning (58-59bp) and the middle (517-518bp) of *tuf* gene that comprise highly variable regions among those regions that allow species differentiation. The *tuf1* and *tuf2* amplicons were adapted to Illumina Miseq300 platform that allows straightforward analysis of amplicon sequencing results. Our primers incorporate traits that make them applicable universally in the *Bacillus* genus where highly variable regions allow for species identification and sequence in high-throughput contexts. While all amplicons were assigned to NCBI database for parallel comparison, a customized *tuf* gene database could potentially improve the resolution of species identification [37]. For instance, retrieving gene sequences encoding members exclusively belonging to the protein family TIGR00485 that translate elongation factor Tu (EF-Tu) gene and use corresponding nucleotide sequences as our database [34]. With the rapid development of long-read sequencing and the study of *Bacillus* evolutionary history, the *tuf* database can be expanded, and a more accurate phylogenomics of *Bacillus* can be established, which could potentially lead to strain-level differentiation in the future. To resolve bacteria community at species level, efforts have been made to increase the identity threshold from 97% to 99%, replace identity based clustering operation taxonomic unit (OTU) with denoising methods amplicon sequence variants (ASV), and create more advanced analysis algorithms [71, 72]. Still, it is challenging to resolve *Bacillus* community composition in natural soil samples by V3-V4 amplicon sequencing. The *tuf* amplicon sequencing methodology reveals the composition of *Bacillus* genera in natural soil samples with highly similar results across biological replicates. To further broaden the use of *tuf* amplicon tool in natural environments, additional experiments that apply metagenome sequencing on same samples to evaluate the performance are necessary. Future applications of the *tuf* primer pairs on different samples from diverse environments, such as rhizosphere, sediments will further examine the sensitivity of *tuf* amplicon methodology.

Nowadays, several *Bacillus* species have been exploited as biofertilizers, biofungicides, or biopesticides. The development of *Bacillus* amplicon sequencing tool (BAST) is essential for agriculture in several aspects. Firstly, evaluating the performance of PGPRs in terms of coexistence, anti-interference, and stabilization is crucial where BAST provides a way to track and identify the species in field. Secondly, accurate identification aids studying plant-microbiome interactions. For instance, with the knowledge that *B. velezensis* produces various lipopeptides, including surfactin, fengycin, and iturin, we can speculate a potentially beneficial interaction with the plant host. Furthermore, for studies involving isolation of *Bacillus* genus in harsh environments, our amplicon sequencing tool would identify the species that alleviated abiotic stress without characterizing the physiological, biochemical properties. In addition, profiling the composition of all stress-tolerant *Bacillus* populations can help identify the best biocontrol performers.

In summary, we designed novel primers and compared with previously documented primers for identification of *Bacilli* at species level. We have exploited our *tuf* gene-targeting primers to accurately classify *Bacillus* on the species level and applied for high-throughput sequencing as a complementary tool in addition to standard 16S amplicon sequencing. The *Bacillus* amplicon sequencing tool (BAST) could be potential applied on tracking bio-inoculant activeness in field, understanding ecological roles of *Bacillus* species in plants, and guiding exploration of bioactive strains in field.

## Acknowledgements

X.X. was supported by a Chinese Scholarship Council fellowship. G.M. was supported by the Lendület-Programme of the Hungarian Academy of Sciences (LP2020-5/2020). This project was supported by the Danish National Research Foundation (DNRF137) for the Center for Microbial Secondary Metabolites and the Novo Nordisk Foundation within the INTERACT project of the Collaborative Crop Resiliency Program (NNF19SA0059360).

## Notes

### Competing Interest Statement

The authors have declared no competing interest.

