## Supplementary material for "Enhanced specificity of *Bacillus* metataxonomics using a *tuf*-targeted amplicon sequencing approach": Fig S1 to S4 and Table S1 to S3

**Dataset S1** List of *Bacillus* genomes downloaded from NCBI in April 2022

**Dataset S2** Suggested primers for *tuf*, *gyrA*, and *rpoB* loci

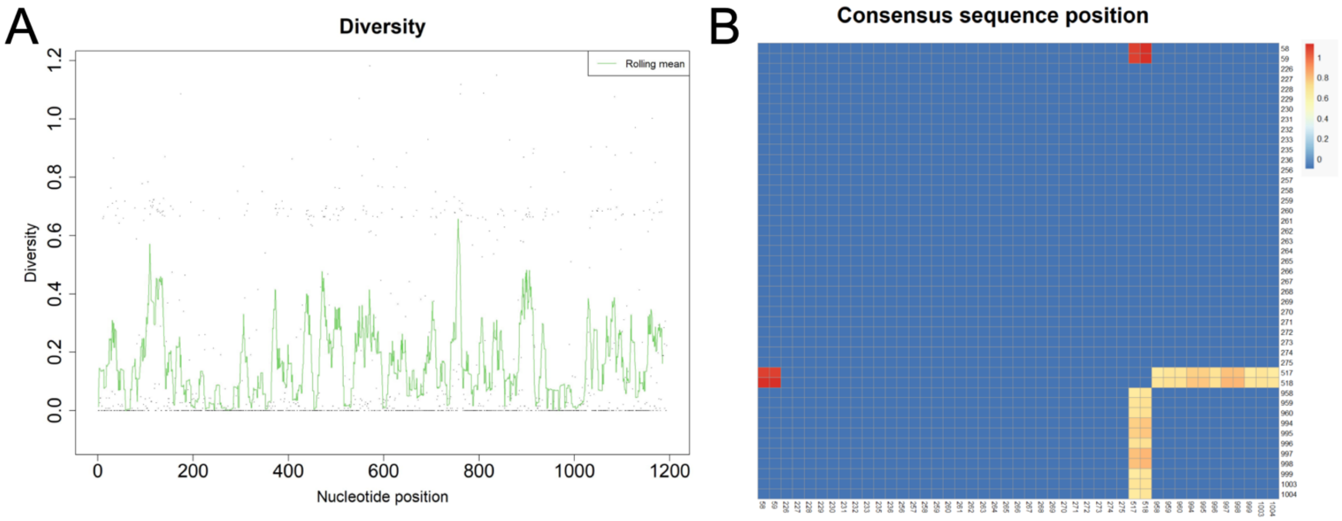


**Fig. S1** Selecting primer positions in the *tuf* gene. **A** Nucleotide entropy profile in a alignment of tuf genes derived from *Bacillus* genome collection. **B** Highly scoring primer position corresponding to the entropy profile of the *tuf* gene alignment.

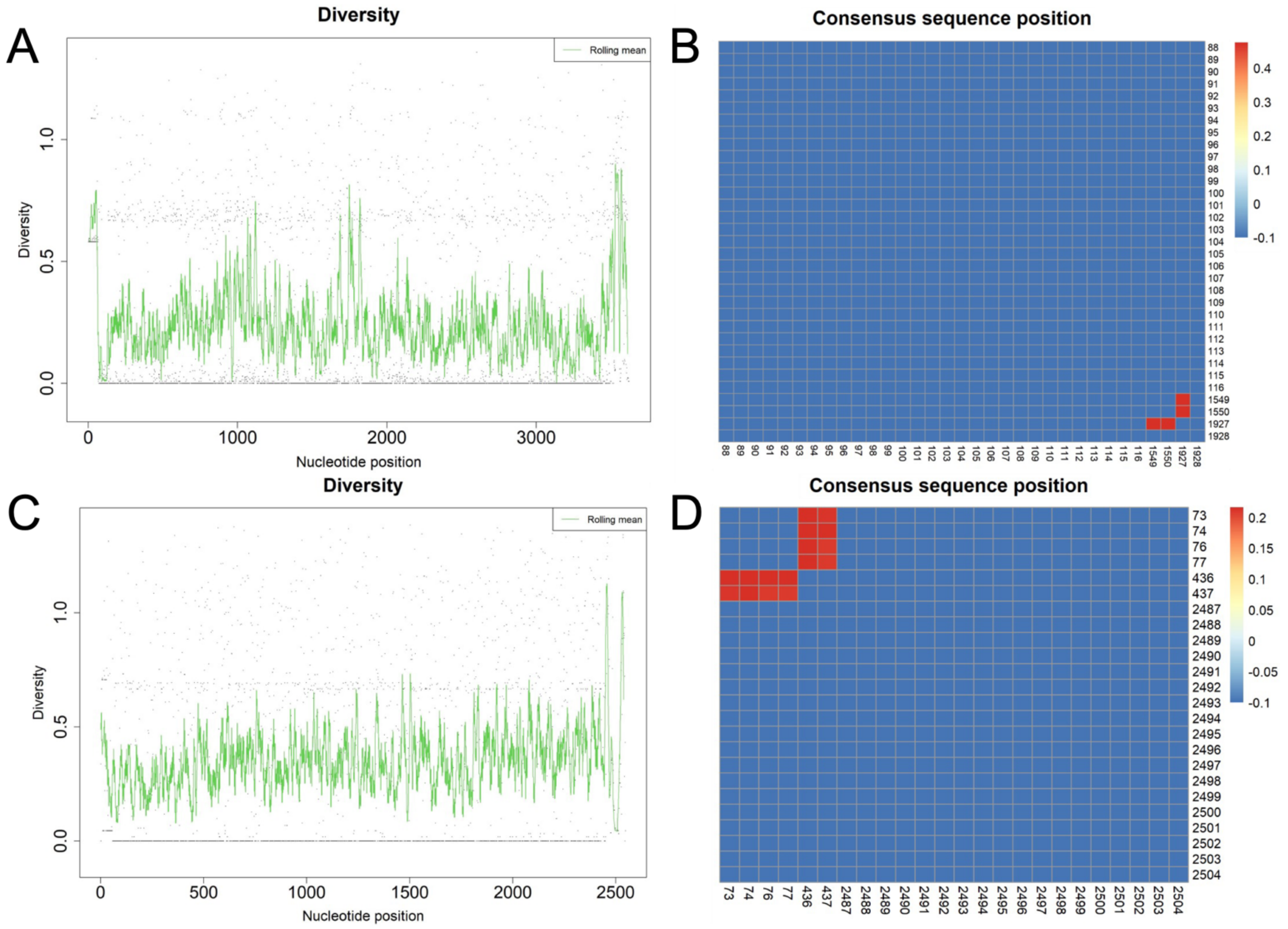


**Fig. S2** Selecting primer positions in the *rpoB* and *gyrA* gene.**A** Nucleotide entropy profile in *rpoB* gene alignment. **B** Highly scoring primer position corresponding to the entropy profile of the *rpoB* gene alignment. **C** Nucleotide entropy profile in *gyrA* gene alignment. **D** Highly scoring primer position corresponding to the entropy profile of the *gyrA* gene alignment.

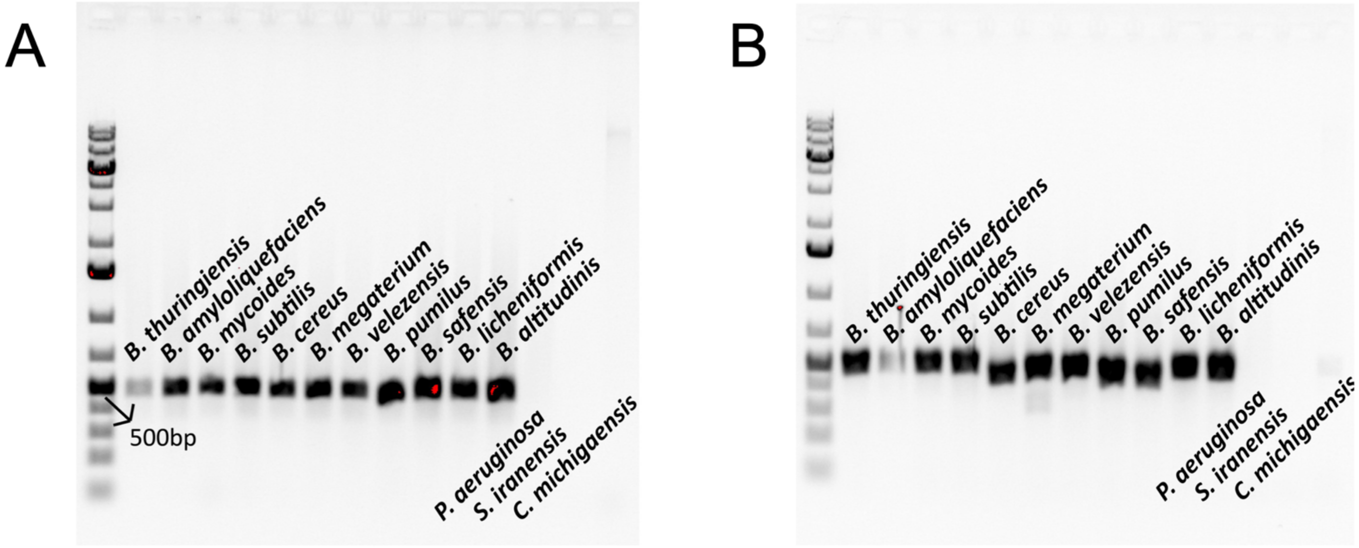


**Fig. S3** Agarose gel electrophoresis of *tuf* gene PCR products amplified from members of Bac-DNAmix using *tuf1* and *tuf2* primers. The last three lanes are non-*Bacillus* strains *Pseudomonas aeruginosa, Streptomyces iranensis, and Clavibacter michiganensis* used as negative controls. Image has been contrast enhanced for better visibility of any band produced. The size marker is Gene ruler 1 kb DNA Ladder (Thermo Fisher Scientific Catalog #SM0311).

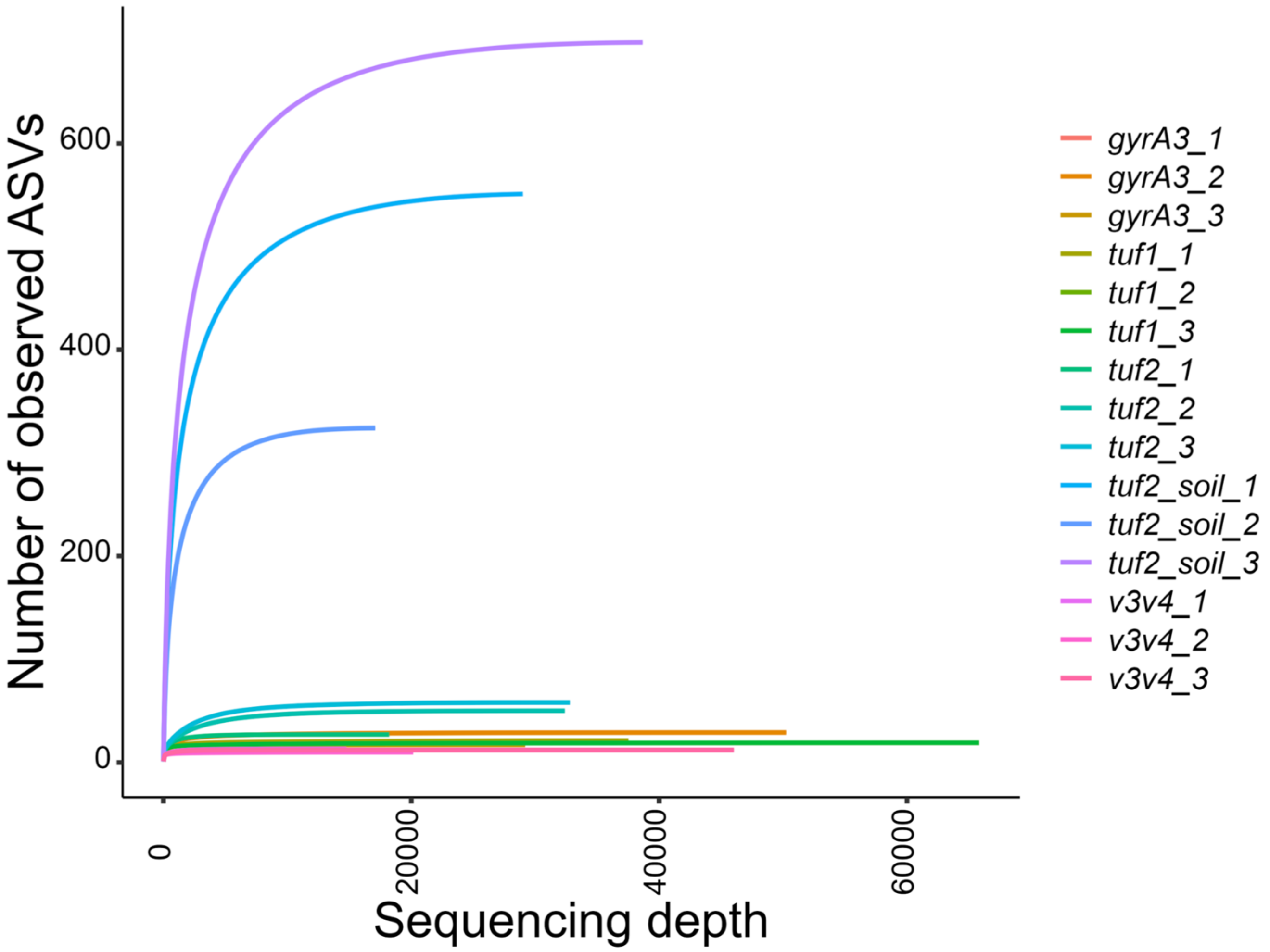


**Fig. S4** Rarefaction curve for each sample and replicates.

**Table S1** List of barcodes tagged on primers for Illumina sequencing of amplicons of the *Bacillus tuf* gene.

| Sample | Barcode |
|---|---|
| tuf1_1 | TTTTAATC |
| tuf1_2 | ATAATTAG |
| tuf1_3 | ACCAAATT |
| tuf2_1 | CTTATCAA |
| tuf2_2 | TGATCATT |
| tuf2_3 | AGAATCTA |
| gyrA3_1 | TCAAGAAA |
| gyrA3_2 | ATCGAAAT |
| gyrA3_13 | ACATTTAC |
| tuf2_soil_1 | TAGAAAAC |
| tuf2_soil_2 | TTATCACC |
| tuf2_soil_3 | AATAGGGT |
| v3v4_1 | GAGAGGGA |
| v3v4_2 | ATCCCGGT |
| v3v4_3 | GAGCTCCT |

**Table S2:** List of non-*Bacillus* genomes tested in silico to validate the specificity of *Bacillus* amplicon primers.

| Phyla/Division | Species | Strain | RefSeq accession |
|---|---|---|---|
| Actinobacteria | *Bifidobacterium bifidum* | S6 | GCF_003390735.1 |
| Actinobacteria | *Clavibacter michiganensis* | UF1 | GCF_009739655.1 |
| Actinobacteria | *Mycobacterium tuberculosis* | HN-506 | GCF_002357975.1 |
| Actinobacteria | *Micrococcus luteus* | CW.Ay | GCF_019890915.1 |
| Actinobacteria | *Rhodococcus globerulus* | D757 | GCF_019334125.1 |
| Actinobacteria | *Streptomyces coelicolor* | A3(2) | GCF_000203835.1 |
| Actinobacteria | *Streptomyces iranensis* | DSM 41954 | GCF_017874715.1 |
| Alphaproteobacteria | *Azospirillum brasilense* | Az39 | GCF_000632475.1 |
| Alphaproteobacteria | *Agrobacterium tumefaciens* | Ach5 | GCF_000971565.1 |
| Alphaproteobacteria | *Bradyrhizobium diazoefficiens* | 110spc4 | GCF_004359355.1 |
| Ascomycota | *Alternaria alternate* | SRC1lrK2f | GCF_001642055.1 |
| Ascomycota | *Fusarium oxysporum* | NRRL 32931 | GCF_000271745.1 |
| Ascomycota | *Saccharomyces cerevisiae* | S288C | GCF_000146045.2 |
| Betaproteobacteria | *Achromobacter xylosoxidans* | AX1 | GCF_008432465.1 |
| Betaproteobacteria | *Achromobacter xylosoxidans* | MN001 | GCF_001051055.1 |
| Betaproteobacteria | *Bordetella pertussis* | H640 | GCF_004008975.1 |
| Betaproteobacteria | *Neisseria meningitidis* | 7-Nov | GCF_008330805.1 |
| Deltaproteobacteria | *Myxococcus xanthus* | DK 1622 | GCF_000012685.1 |
| Firmicutes | *Aneurinibacillus migulanus* | DSM 2895 | GCF_001274715.1 |
| Firmicutes | *Clostridium acetobutylicum* | ATCC 824 | GCF_000008765.1 |
| Firmicutes | *Lactococcus lactis* | LAC460 | GCF_020463755.1 |
| Firmicutes | *Paenibacillus amylolyticus* | FSL H7-0692 | GCF_001956035.1 |
| Firmicutes | *Paenibacillus graminis* | DSM 15220 | GCF_000758705.1 |
| Firmicutes | *Paenibacillus polymyxa* | ZF129 | GCF_006274405.1 |
| Firmicutes | *Staphylococcus aureus* | NCTC 8325 | GCF_000013425.1 |
| Firmicutes | *Staphylococcus epidermidis* | NIHLM061 | GCF_000276445.1 |
| Flavobacteriia | *Chryseobacterium sp.* | D764 | GCF_019317325.1 |
| Gammaproteobacteria | *Acinetobacter baumannii* | K09-14 | GCF_008632635.1 |
| Gammaproteobacteria | *Enterobacter hormaechei subsp. xiangfangensis* | ND10 | GCF_000784035.1 |
| Gammaproteobacteria | *Legionella pneumophila* | C9_S | GCF_001753085.1 |
| Gammaproteobacteria | *Pseudomonas aeruginosa* | PAO1 | GCF_000006765.1 |
| Gammaproteobacteria | *Pseudomonas fluorescens* | ATCC 13525 | GCF_900215245.1 |
| Gammaproteobacteria | *Pseudomonas koreensis* | LMG21318 | GCF_900101415.1 |
| Gammaproteobacteria | *Pseudomonas moraviensis* | TYU6 | GCF_002287825.1 |
| Gammaproteobacteria | *Pseudomonas protegens* | CHA0 | GCF_900560965.1 |
| Gammaproteobacteria | *Pseudomonas stutzeri* | F2a | GCF_019704535.1 |
| Gammaproteobacteria | *Stenotrophomonas indicatrix* | D763 | GCF_019285675.1 |
| Gammaproteobacteria | *Vibrio parahaemolyticus* | O3:K6 substr. RIMD 2210633 | GCF_000196095.1 |
| Nematoda | *Caenorhabditis elegans* | Bristol N2 | GCF_000002985.6 |
| Sphingobacteria | *Sphingobacterium sp.* | B29 | GCF_001952815.1 |
| Sphingobacteria | *Pedobacter sp.* | D749 | GCF_019317285.1 |
